# Habitat specialization shapes the evolution of transcriptional responses to hypoxia

**DOI:** 10.64898/2026.05.05.723024

**Authors:** Maribet Gamboa, Mauren Vergara, Emily Winter, Brian K Hand, Gordon Luikart, Jack A Stanford, Rachel L Malison

## Abstract

Oxygen limitation is a widespread environmental constraint that shapes physiological and evolutionary responses across ecosystems. A central unresolved question is whether tolerance to hypoxia reflects generalized stress responses or coordinated regulatory strategies shaped by long-term environmental exposure. Here, we use comparative transcriptomic analyses to examine gene expression responses to low oxygen in two aquifer-dwelling stoneflies (*Isocapnia* sp. and *Paraperla frontalis*) and one benthic species (*Sweltsa* sp.) under controlled conditions. Time-series analysis in *Isocapnia* sp. revealed a multi-phase transcriptional response involving early regulatory activation, metabolic reorganization, and late-stage cellular stabilization. Across aquifer taxa, hypoxia was associated with downregulation of energy-demanding processes and upregulation of pathways related to oxidative stress mitigation, metabolite transport, and protein folding, consistent with coordinated cellular adjustment to oxygen limitation. In contrast, the river benthic species exhibited transcriptional profiles dominated by neural signaling, ion channel activity, and structural remodeling, which are patterns consistent with acute physiological stress rather than coordinated regulation. Despite these differences, all taxa showed modulation of ion transport and calcium signaling pathways, suggesting conserved mechanisms of hypoxia sensing. Together, these results indicate that transcriptional responses to hypoxia differ systematically with habitat and are consistent with the evolution of distinct regulatory strategies in chronically hypoxic environments.

**Significant statement:** Oxygen limitation is a common environmental challenge that affects organisms across aquatic and terrestrial ecosystems, yet the mechanisms by which species cope with low oxygen remain incompletely understood. A key question is whether tolerance to hypoxia reflects common stress responses or the evolution of coordinated metabolic regulatory strategies under chronic exposure. By comparing gene expression responses in closely related aquatic insects from oxygen-variable underground aquifers and oxygen-rich river habitats, we show that species that evolved under persistent hypoxia exhibit transcriptional patterns consistent with energy conservation and cellular stabilization, whereas those experiencing hypoxia as a transient stress display signature of physiological disruption. These findings highlight fundamental differences between evolutionarily adaptive and acute stress-driven responses to environmental change and provide insight into how organisms may respond to increasing hypoxia under global change.

## Introduction

Oxygen limitation is a fundamental constraint across aquatic and terrestrial ecosystems, shaping physiological performance, ecological interactions, and evolutionary trajectories. A central question in environmental biology is how organisms cope with hypoxia (1, 2), particularly whether tolerance arises from short-term modulation of conserved stress-response pathways or from the evolution of specialized regulatory strategies under chronic exposure. While acute hypoxia typically triggers generalized stress responses (including disruption of cellular homeostasis, activation of ion transport, and metabolic imbalance; (3; 4) organisms inhabiting persistently low-oxygen environments may exhibit fundamentally different, coordinated mechanisms that have evolved to enable long-term survival.

Understanding this distinction between acute stress responses and evolved tolerance is critical for predicting how species respond to environmental change, including the increasing prevalence of hypoxia driven by climate change (warming) and anthropogenic impacts. However, most mechanistic studies of hypoxia have focused on model organisms or short-term exposures (5), limiting our ability to infer how long-term environmental conditions shape transcriptional regulation and adaptive strategies in natural systems.

Alluvial aquifers provide a powerful natural context to address this question. These subsurface habitats are characterized by variable oxygen conditions, including zones of hypoxia and anoxia (6,9), limited organic carbon, and strong physicochemical gradients (7, 8), yet they support abundant and specialized invertebrate communities. Among these, aquifer-dwelling stoneflies (Plecoptera) exhibit remarkable tolerance to hypoxia and anoxia (10–13), maintaining activity and survival under conditions that are lethal to closely related benthic river taxa. Previous work has shown that aquifer species can sustain aerobic metabolism at low oxygen concentrations and tolerate prolonged anoxia (14–16), suggesting the presence of evolved physiological mechanisms (15). However, the molecular basis of this tolerance (and whether it reflects modified stress responses or distinct regulatory profiles) remains unresolved.

Insects generally respond to hypoxia through conserved pathways involving metabolic suppression (17), oxidative stress regulation (18), and cellular homeostasis, often mediated by hypoxia-inducible factors and associated signaling networks (18–20). Yet, a key unresolved question in evolutionary physiology is whether tolerance to chronic hypoxia arises from modifications of conserved stress-response pathways or from the evolution of distinct regulatory profile. Natural systems that contrast long-term exposure to hypoxia with transient exposure provide an opportunity to disentangle these alternatives.

Here, we use comparative transcriptomics to test how organisms respond to chronic versus acute hypoxia in a natural evolutionary context. We analyze gene expression under controlled low-oxygen conditions in two aquifer taxa (*Isocapnia* sp. and *Paraperla frontalis*) and one benthic taxon (*Sweltsa* sp.), integrating time-series and differential expression approaches to identify both shared and habitat-specific responses. We hypothesize that aquifer stoneflies exhibit coordinated pathway of transcriptional profile consistent with evolved tolerance to chronic hypoxia, characterized by metabolic reorganization, energy conservation, and enhanced cellular maintenance, whereas benthic taxa exhibit transcriptional profiles characteristic of stress responses indicative of acute physiological disruption. By contrasting these systems, we aim to investigate whether long-term exposure to hypoxia leads to the emergence of distinct regulatory strategies, providing broader insight into the mechanisms by which organisms adapt to extreme environments.

## Results

### Aquifer species exhibit coordinated transcriptional responses to hypoxia

To evaluate transcriptional responses to oxygen limitation, we conducted a time-series analysis in *Isocapnia* sp. Temporal clustering revealed nine distinct expression trajectories, indicating a highly coordinated response to hypoxia (Fig. 1A). These trajectories define three major phases: early regulatory activation (clusters 1–4), intermediate metabolic adjustment (clusters 5–7), and late-stage stabilization (clusters 8–9).

**Figure 1.**
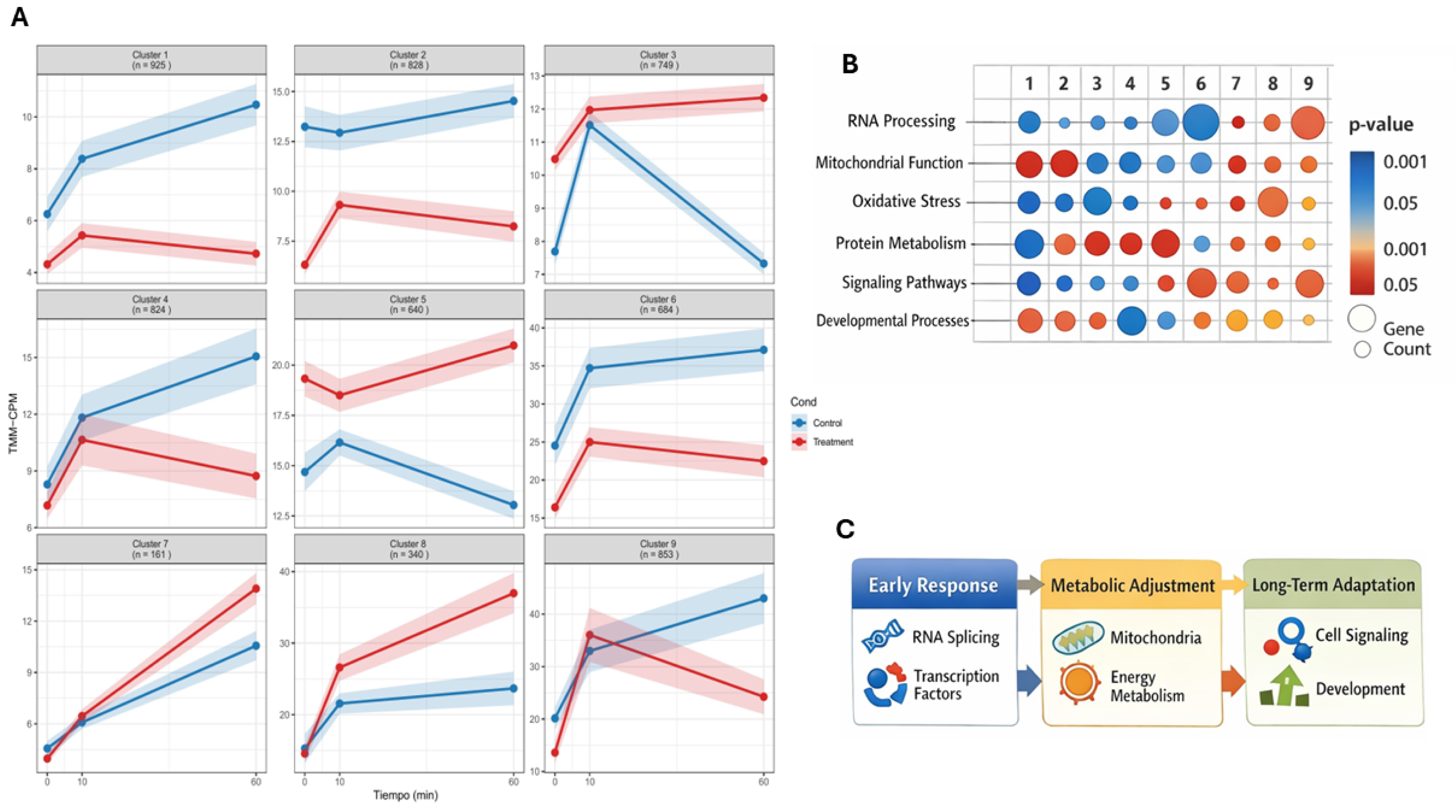
Temporal gene expression dynamics and functional enrichment under hypoxia treatment of *Isocapnia* sp. **(A)** Time-series clustering of differentially expressed genes across 0, 10, and 60 min. Nine clusters were identified based on similar expression trajectories (TMM-normalized CPM). Lines show mean expression profiles with shaded areas representing standard error. Blue indicates control, and red indicates treatment. The number of genes per cluster is shown in each panel. **(B)** Functional enrichment of gene clusters. Bubble plots summarize enriched GO and KEGG categories. Circle size represents gene count, and color indicates enrichment significance (p-value). **(C)** Conceptual model of the transcriptional response in three phases.

Functional enrichment analyses revealed hierarchical organization of biological processes across these phases (Fig. 1B). Early responses were dominated by transcriptional regulation, RNA processing, and signal transduction (Supplementary Fig. 1A-D), indicating rapid cellular sensing of oxygen depletion. These were followed by enrichment of pathways associated with mitochondrial function, energy metabolism, oxidative processes, and metabolite transport (Supplementary Fig. 1E-G), consistent with evolved metabolic reprogramming. Late-stage responses involved signaling pathways, developmental processes, and intercellular communication (Supplementary Fig. 1H-I), suggesting sustained physiological adjustment under prolonged hypoxia. This transcriptional trajectory reveals a coordinated response to oxygen limitation and may reflect *Isocapnia* sp. adaptation to hypoxic environments (Fig. 1C).

Consistent with these temporal dynamics, differential expression analysis identified 749 genes (Supplementary Fig. 3A) significantly affected by hypoxia (FDR < 0.05), with a balanced distribution of upregulated (362) and downregulated (387) transcripts (Supplementary Fig. 3B). Hypoxia was associated with increased expression of genes involved in oxidative stress mitigation, metabolite transport (including ABC transporters and hexamerins), and protein folding, whereas genes associated with cellular organization and regulatory processes were downregulated (Supplementary Table 2, Supplementary Fig. 2). Together, these results indicate a coordinated transcriptional program that transitions from rapid sensing to metabolic adjustment and ultimately to cellular stabilization under sustained hypoxia.

### Aquifer taxa share differences in oxidative stress regulation

Strong transcriptional responses to hypoxia were also observed in *P. frontalis*. Differential expression and multivariate analyses revealed clear separation between control and treatment groups, indicating robust transcriptional shifts associated with oxygen limitation (Fig. 2A–C). Functional enrichment analyses identified pathways (Fig. 2D) related to cell cycle regulation, DNA replication and repair, carbon metabolism, and glyoxylate metabolism, as well as processes associated with cellular maintenance and structural regulation (Supplementary Fig. 4A-C, Supplementary Table 3).

**Figure 2.**
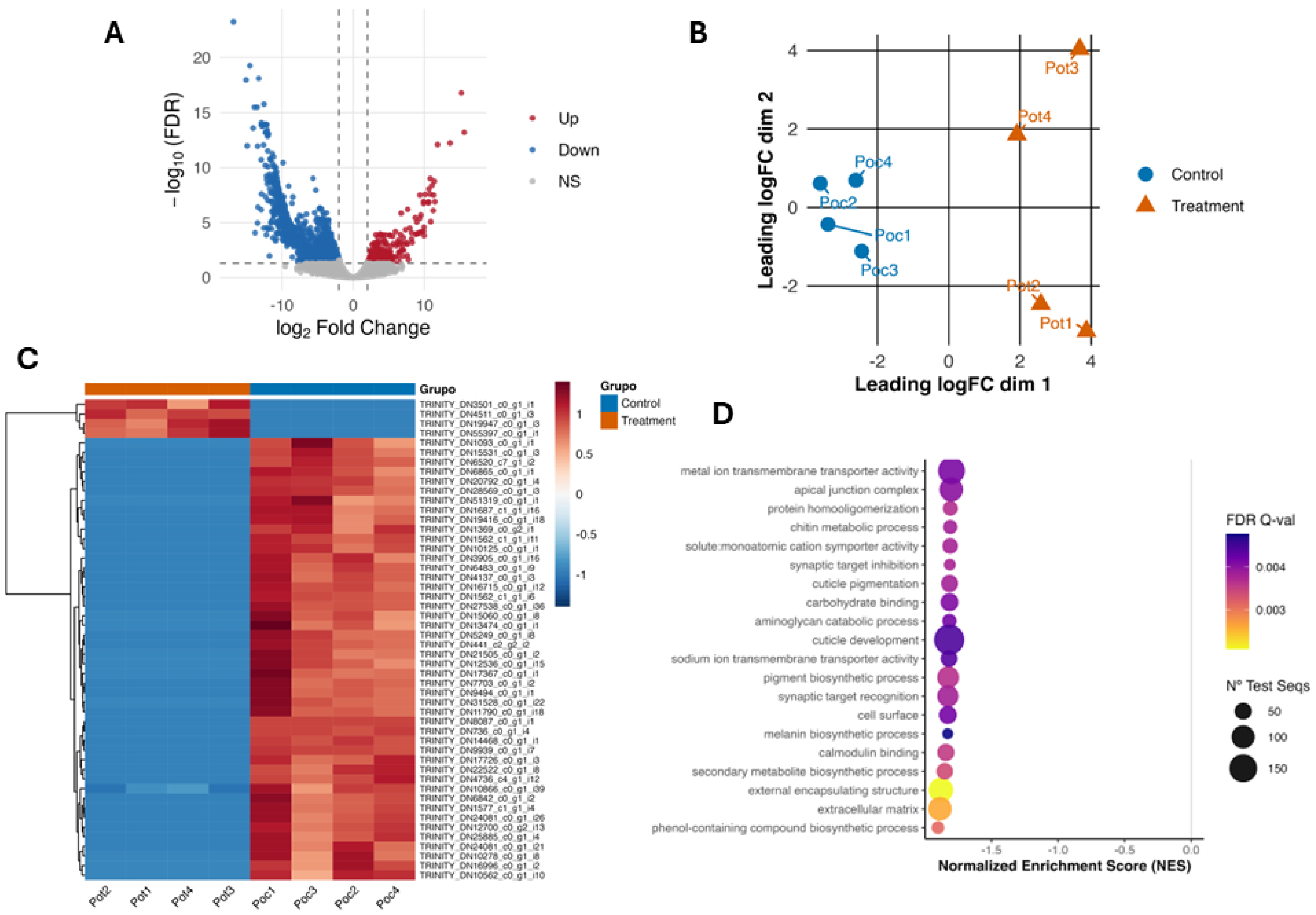
Differential gene expression and functional enrichment under hypoxia treatment of *Paraperla frontalis*. **(A)** Volcano plot showing differentially expressed genes between treatment and control. Red and blue represent significantly upregulated and downregulated genes, respectively, while grey points indicate non-significant genes. **(B)** Multidimensional scaling (MDS) plot based on leading log fold-change distances among samples, showing separation between control (blue) and treatment (orange) groups. **(C)** Heatmap of significantly differentially expressed genes across samples. Colors represent scaled expression levels, with hierarchical clustering highlighting distinct transcriptional patterns between control and treatment conditions. **(D)** GSEA pathway enrichment analysis of the top 20 differentially expressed genes. Bubble size indicates the number of genes associated with each pathway, while color represents the normalized enrichment score (NES).

Despite these shared responses, comparison with *Isocapnia* sp. revealed important differences in stress-response strategies. Both aquifer taxa exhibited downregulation of energy-expensive processes, including secondary metabolism and extracellular matrix organization, consistent with energy conservation under hypoxia. However, *Isocapnia* sp. showed stronger enrichment of oxidative stress-related pathways and chaperone activity, suggesting a greater capacity for managing reactive oxygen species (ROS), chemically reactive molecules that can damage cellular components. This contrast indicates that while energy conservation represents a common strategy among aquifer taxa, the magnitude and nature of oxidative stress responses may differentiate levels of hypoxia tolerance, possibly suggesting a more effective physiological strategy for coping with low-oxygen conditions.

### Benthic species exhibit acute physiological stress

In contrast to aquifer taxa, the benthic river species *Sweltsa* sp. displayed transcriptional responses consistent with acute physiological stress. Differential expression analyses revealed substantial transcriptional changes under hypoxia, with clear separation between control and treatment groups (Fig. 3A–C).

**Figure 3.**
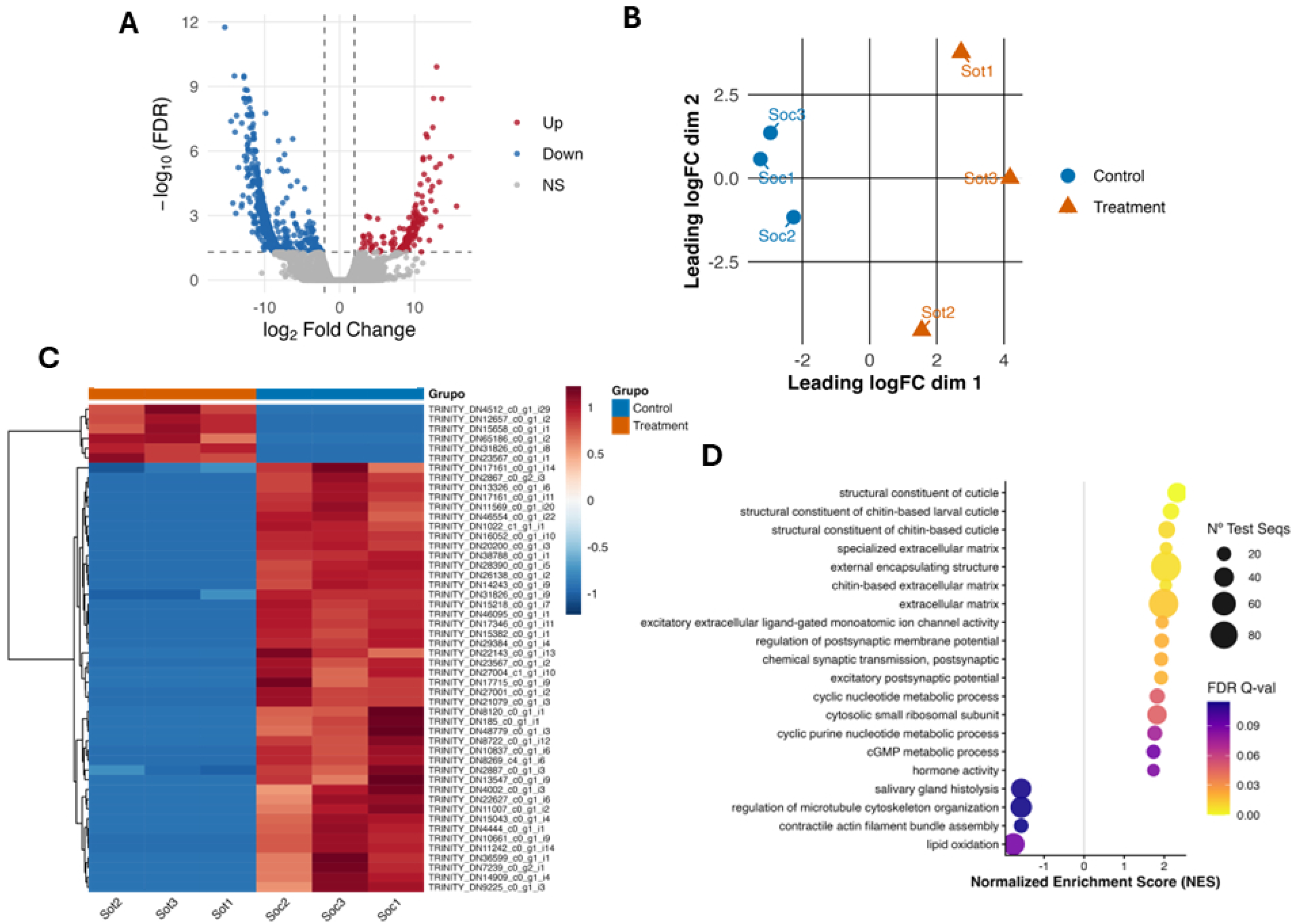
Differential gene expression and functional enrichment under hypoxia treatment of *Sweltsa* sp. **(A)** Volcano plot showing differentially expressed genes between treatment and control. Red and blue represent significantly upregulated and downregulated genes, respectively, while grey points indicate non-significant genes. **(B)** Multidimensional scaling (MDS) plot based on leading log fold-change distances among samples, showing separation between control (blue) and treatment (orange) groups. **(C)** Heatmap of significantly differentially expressed genes across samples. Colors represent scaled expression levels, with hierarchical clustering highlighting distinct transcriptional patterns between control and treatment conditions. **(D)** GSEA pathway enrichment analysis of the top 20 differentially expressed genes. Bubble size indicates the number of genes associated with each pathway, while color represents the normalized enrichment score (NES).

Functional enrichment analyses identified pathways associated with neural signaling, ion channel activity, structural remodeling, and amino acid metabolism (Fig. 3D). Notably, genes related to cuticle structure, ligand-gated ion channels, and synaptic signaling were strongly represented (Supplementary Table 4), suggesting activation of pathways involved in maintaining cellular integrity and signaling under stress. Additional responses included modulation of lipid metabolism and proteolysis (Supplementary Fig. 5), indicating metabolic imbalance rather than coordinated energy regulation.

Together, these patterns indicate that *Sweltsa* sp. responds to hypoxia through generalized stress responses, characterized by structural and signaling disruption, rather than through coordinated regulatory programs that apparently evolved in the aquifer taxa.

### Comparative analysis reveals divergent transcriptional strategies between chronic and acute hypoxia exposure

Comparative analysis across species revealed distinct transcriptional strategies between aquifer and the benthic taxa (Table 1). Aquifer species (*Isocapnia* sp. and *P. frontalis*) exhibited consistent downregulation of processes associated with high energetic demand, including secondary metabolite biosynthesis, extracellular matrix organization, protein complex assembly, and pigmentation pathways. These patterns are consistent with a coordinated reduction of non-essential processes, indicative of energy conservation under chronic hypoxia.

**Table 1.**
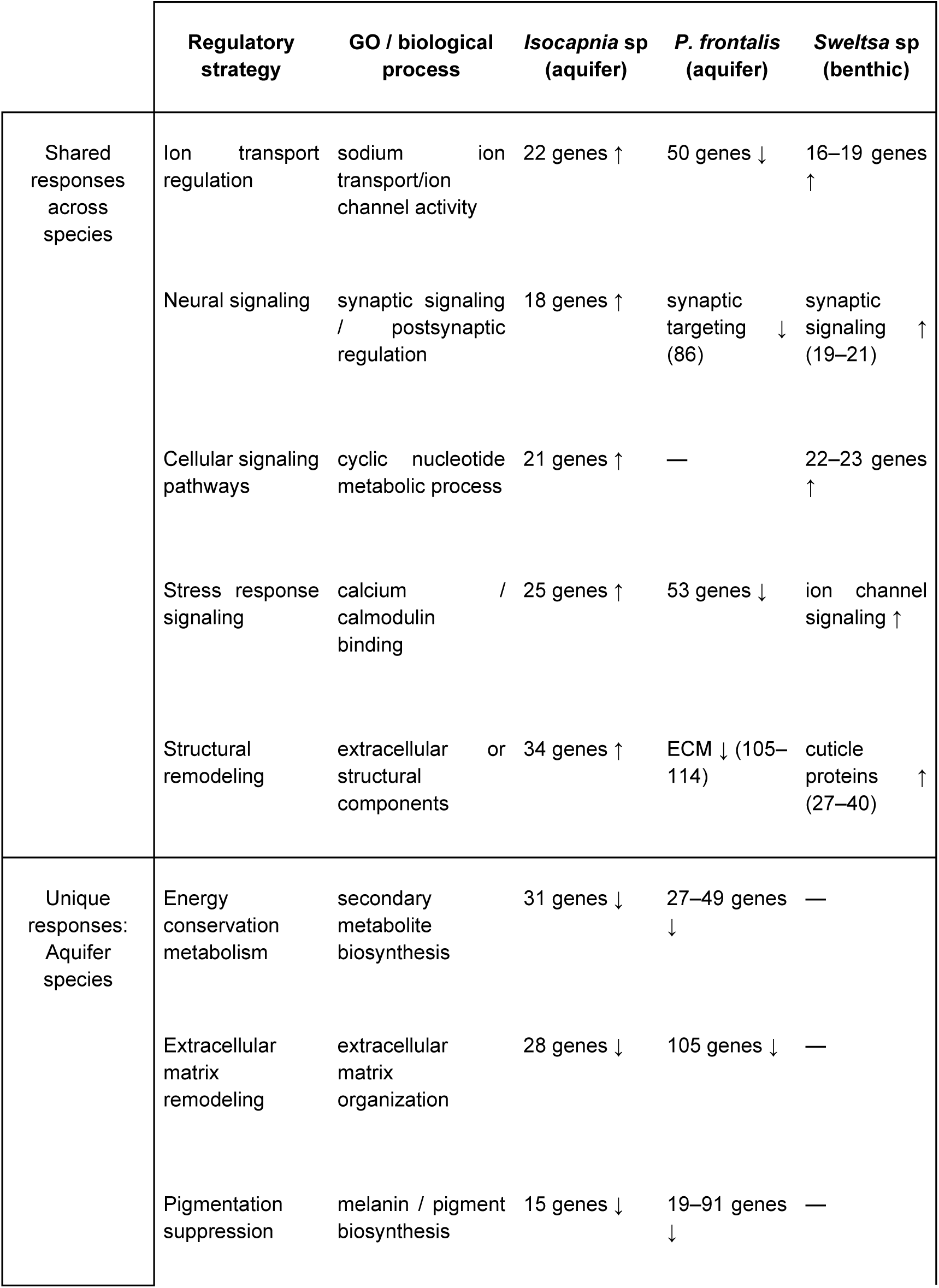
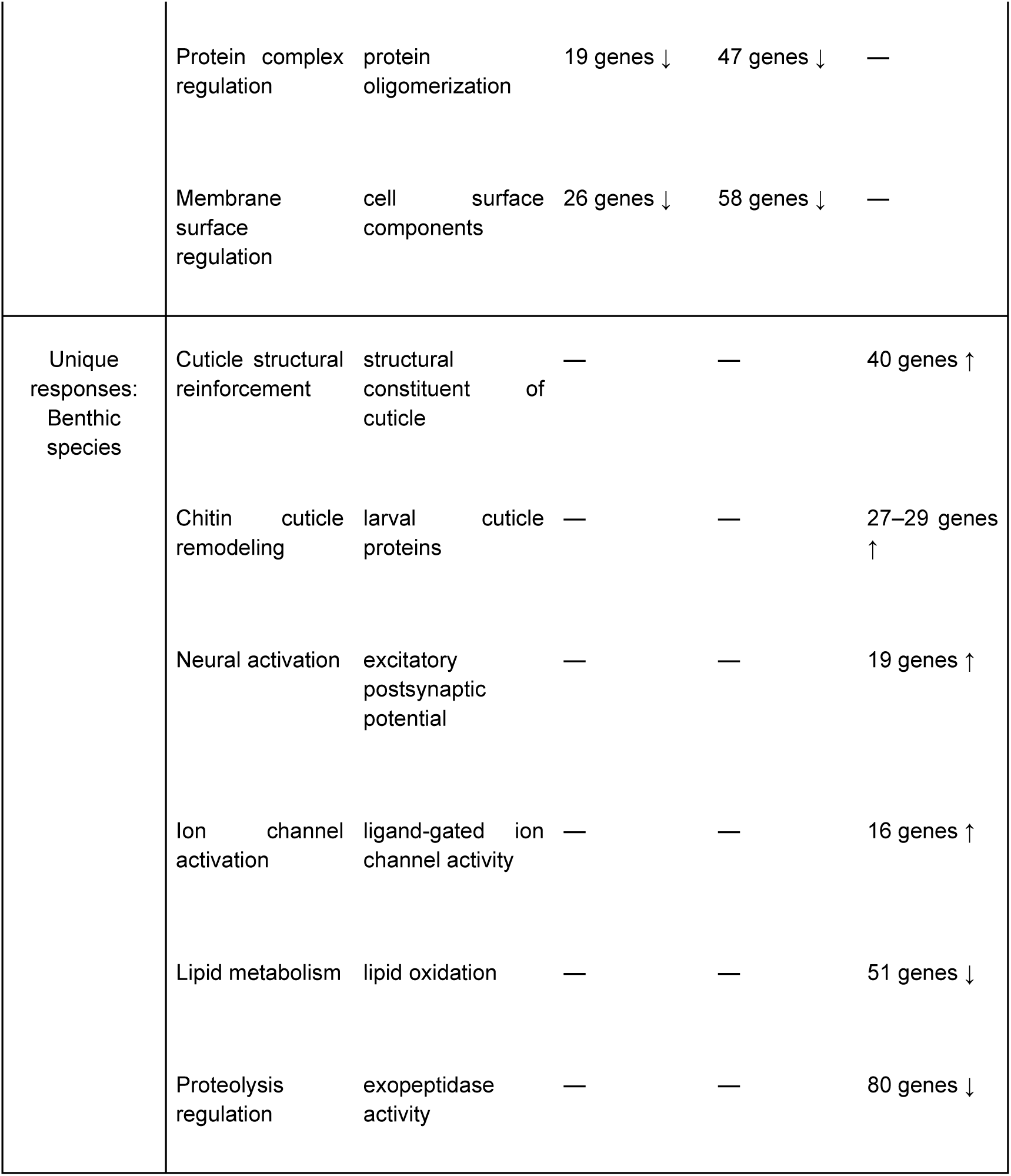
Comparative ortholog functional responses to hypoxia treatment of the top 5 genes.

In contrast, the benthic species *Sweltsa* sp. displayed unique enrichment of pathways associated with cuticle reinforcement, neural activation, and ion channel signaling, reflecting physiological stress responses rather than targeted metabolic adjustment. These responses were accompanied by downregulation of lipid metabolism and proteolysis-related pathways, further indicating disruption of metabolic homeostasis.

Despite these differences, several responses were conserved across all taxa. Ion transport and channel activity were broadly modulated and signaling pathways involving cyclic nucleotides and calcium were consistently enriched, indicating shared mechanisms of hypoxia sensing. However, the downstream transcriptional responses diverged markedly, with aquifer taxa exhibiting coordinated metabolic reorganization and cellular maintenance, whereas the benthic species displayed acute low oxygen stress transcriptional disruption.

Collectively, these results support a model in which chronic hypoxia favors the emergence of coordinated transcriptional programs that prioritize energy conservation and cellular stability, whereas acute hypoxia elicits transcriptional profiles characteristic of generalized stress responses (i.e., ion channel activation, lipid metabolism). These differences suggest that habitat specialization strongly influences molecular responses to oxygen limitation.

## Discussion

Hypoxia regulates genes and enzymes involved in protection against low dissolved oxygen in several taxa (19), including insects (21–22). This study provides novel insights into the molecular mechanisms underlying hypoxia tolerance in aquifer stoneflies by integrating time-series and comparative transcriptomic analyses. Our results support a general model in which chronic exposure to hypoxia favors coordinated transcriptional programs that prioritize energy conservation and cellular stability, whereas acute hypoxia elicits generalized stress responses characterized by structural and signaling disruption.

### A coordinated, multi-phase response underlies hypoxia tolerance in aquifer species

The time-series analysis of *Isocapnia* sp. demonstrated that hypoxia tolerance is not mediated by a single pathway but by a temporally structured, multi-phase transcriptional response, as observed previously in different taxa (23). Early activation of genes involved in transcriptional regulation, RNA processing, and signaling suggests rapid cellular sensing of oxygen depletion and initiation of regulatory cascades. This is followed by intermediate responses associated with mitochondrial function, energy metabolism, and oxidative processes, and finally by late-stage processes linked to cellular communication and long-term physiological adjustment.

Such temporal structuring may represent adaptive plasticity to fluctuating oxygen conditions, where early regulatory control enables downstream metabolic reprogramming. The observed cascade (from signaling to metabolic adjustment and eventual stabilization) aligns with models of hypoxia response described in other insects, where transcriptional regulation precedes metabolic suppression and stress mitigation (19, 24). Importantly, the enrichment of pathways associated with detoxification, protein folding, and oxidative stress protection suggests that *Isocapnia* sp. is not only tolerating hypoxia but actively mitigate the cellular damage associated with hypoxia-reoxygenation cycles, as previously observed in hypoxia-induced fish (25) and bivalves (26).

### Differential metabolic strategies distinguish aquifer taxa

Comparisons between aquifer species revealed both shared and species-specific strategies. Both *Isocapnia* sp. and *P. frontalis* showed downregulation of energy-expensive processes such as secondary metabolite biosynthesis, extracellular matrix organization, and protein complex assembly. This pattern supports the hypothesis that aquifer species employ energy-conservation strategies under hypoxia, reducing non-essential cellular functions to maintain homeostasis. This downregulation aligns with reports from other taxa in which hypoxia limits processes in favor of stress tolerance mechanisms (e.g., plants, 27; vertebrates, 28; insects, 29).

However, a key difference between aquifer taxa lies in the magnitude and nature of oxidative stress responses. *Isocapnia* sp. exhibited stronger enrichment of oxidative stress-related genes and chaperone activity compared to *P. frontalis*, suggesting a more robust capacity to manage reactive oxygen species (ROS). This difference is particularly relevant given that hypoxia and subsequent reoxygenation can lead to oxidative damage (30). The enhanced oxidative stress response in *Isocapnia* sp. may explain its superior tolerance to prolonged anoxia observed in previous physiological studies (15), indicating that ROS management is a key component of extreme hypoxia tolerance.

Additionally, the upregulation of transport-related genes (e.g., ABC transporters) and storage proteins such as hexamerins suggests mechanisms for maintaining cellular resource allocation and metabolite balance under oxygen limitation. These findings are consistent with previous work highlighting the role of hexamerins in energy storage and stress resilience in insects (21, 31).

### Benthic species exhibit stress responses rather than adaptive regulation

In contrast to aquifer taxa, the benthic river species *Sweltsa* sp. displayed transcriptional profiles indicative of acute physiological stress rather than regulatory patterns consistent with evolved tolerance. Enrichment of pathways related to cuticle restructuring, neural activation, and ion channel activity suggests attempts to maintain structural integrity and cellular signaling under unfavorable conditions. However, these responses appear reactive rather than coordinated, anticipatory or preconditioned by chronic exposure, as observed in other organisms (32).

The activation of neural signaling and ligand-gated ion channels may reflect disruption of cellular homeostasis and increased energetic demand (33) rather than efficient adaptation. Similarly, the downregulation of lipid metabolism and proteolysis-related genes suggests metabolic imbalance under hypoxia (34). Together, these patterns support the interpretation that benthic species lack specialized molecular mechanisms for coping with hypoxia and instead exhibit generalized stress responses when exposed to hypoxic conditions.

### Shared responses highlight conserved hypoxia signaling pathways

Despite clear differences, several transcriptional responses were conserved across all taxa, including modulation of ion transport, calcium signaling, and neural pathways. These shared responses likely represent core components of hypoxia sensing in aquatic insects. Calcium signaling, in particular, plays a central role in hypoxia sensing and cellular stress responses, regulating pathways associated with apoptosis (35), metabolism (36), and cytoskeletal dynamics (37) in insects.

The presence of conserved signaling pathways alongside divergent metabolic strategies suggests that while the initial detection of hypoxia may be similar across taxa, downstream responses have diverged according to habitat specialization. This pattern aligns with evolutionary models where conserved regulatory frameworks are modified to produce habitat-specific phenotypes (38–39).

### Evidence for molecular adaptation to hypoxic aquifer environments

Overall, the transcriptional signatures observed in aquifer stoneflies are consistent with long-term adaptation to chronically low-oxygen environments. The combination of energy conservation, enhanced oxidative stress management, and coordinated transcriptional regulation indicates a shift toward efficient cellular maintenance under resource-limited conditions. These transcriptional patterns are consistent with mechanisms shown in previous functional studies of enhanced hypoxia tolerance (30, 40). These findings complement previous ecological and physiological studies demonstrating exceptional hypoxia tolerance in aquifer taxa (15, 16, 21), providing a mechanistic basis for these traits.

Importantly, the lack of reliance on morphological adaptations (e.g., gills or behavioral responses, 41) further supports the hypothesis that hypoxia tolerance in aquifer stoneflies is primarily mediated at the molecular level. The observed transcriptional profile may represent evolved traits that enable sustained activity and survival in environments characterized by extreme and persistent oxygen limitations.

### Limitations and future directions

Some limitations should be considered when interpreting these results. Sample sizes differed among species, particularly for *P. frontalis* and *Sweltsa* sp., which may reduce statistical power for detecting subtle expression changes. Future studies incorporating larger sample sizes, and single-individual transcriptomics, may strengthen inference. Further work should also explore the role of key candidate pathways and genes identified here, particularly those related to oxidative stress, ion transport, and metabolic regulation. Integrating transcriptomic data with physiological and metabolic measurements would provide a more comprehensive understanding of hypoxia tolerance mechanisms. Finally, further studies should observe the differences between nymphs of different instars, as well as adult aquifer stoneflies, to obtain a better understanding of how the regulation of transcriptional organization changes over the course of stonefly development as they prepare to move from aquatic to terrestrial environments.

### Conclusions

This study demonstrates that aquifer stoneflies exhibit distinct, temporally-structured, and coordinated transcriptional responses to hypoxia that differ fundamentally from those of benthic river species. These findings highlight the importance of habitat-driven selection in shaping molecular responses to environmental stress and provide new insight into the mechanisms of hypoxia adaptation in aquatic insects. By linking transcriptional dynamics to ecological specialization, this work advances our understanding of how organisms persist in extreme environments and contributes to broader efforts to integrate functional genomics with ecological and evolutionary theory. Hypoxia is an increasingly prevalent stressor in aquatic ecosystems due to climate change and eutrophication. The distinction between evolutionarily adaptive and acute low oxygen stress transcriptional responses identified here may therefore have broad implications for predicting species resilience under global environmental change.

## Materials and Methods

### Study Area and field sampling

Aquifer and benthic stoneflies were collected from habitats within the Nyack floodplain of the Middle Fork of the Flathead River in northwestern Montana (Supplementary Fig. 6; 48°27’30″ N, 113°50’0” W; see floodplain description in refs. 15, 42). The fifth-order free-flowing Middle Fork has a spring snowmelt hydrograph and flows through the Nyack floodplain (∼10 km in length and ∼2 km wide), forming an expansive “hyporheic zone” when 30% of the river water downwells into the upstream end of the floodplain (42–43). The floodplain is instrumented with an extensive network of wells from which aquifer stoneflies can be sampled (see refs. 15-16).

To measure the effects of low oxygen on gene expression, aquifer and benthic stoneflies were collected on the same day from the Nyack floodplain. Aquifer stoneflies (*Isocapnia* spp. and *Paraperla frontalis*) were collected from the alluvial aquifer by pumping nymphs from wells into a 500 μm net from all wetted depths (2-7 m) using a gas-powered diaphragm pump and hose. Benthic stoneflies were collected from benthic river habitats using a Stanford-Hauer kick net. Once collected, all live individuals were transported back to the Flathead Lake Biological Station (FLBS) laboratory in aerated buckets of cold water. All individuals were held at 5 °C in a refrigerator for 2 days until the experiment.

### Hypoxia variation experiment

Control and treatment baths were held at a constant temperature (∼ 5.5 °C). The control bath was aerated continuously, while the treatment bath was deoxygenated using nitrogen gas. At the beginning of the experiment, between 2-3 stoneflies were placed into individual pouches, and the pouches were placed into normoxic water in both baths at the same time. After this, the nitrogen gas was turned on and the oxygen in the treatment bath was lowered to anoxia (<0.05 mg/L) over the course of 19 minutes. Only individuals of *Isocapnia* sp. were used in the experiment to evaluate responses to temporal oxygen variation. Individuals from both control and treatment groups were sampled at 0 min, 5–10 min, and 1 h of anoxia exposure. Upon removal, each individual (one control and one treatment at the same time) was placed in a labeled cryovial and then flash frozen with liquid nitrogen.

### RNA extraction and sequencing

RNA was extracted from stonefly tissue using NucleoSin RNA kits. Each stonefly was homogenized with a steel bead placed in the tube by running the tissue-lyzer at 30 hz for 1 min. Cells were lysed (using a mixture of 350 μL Buffer RA1 and 3.5 μL ß-mercaptoethanol). The lysate was filtered by NucleoSpin® Filter. The silica membrane was then desalted and the DNA digested (using rDNase Reaction Buffer). The silica membrane was washed and dried and then RNA was eluted with ultrapure water nuclease free. Library preparation (poly(A)-enriched, non-directional) and RNA sequencing were performed by Novogene Corporation Inc. (Sacramento, CA, USA). A total of 95 RNA samples were processed, with sequencing conducted to a depth of approximately 20 million reads per sample.

Several individuals were combined to achieve the required amount of RNA for library construction. Therefore, the downstream analysis included 53 aquifer *Isocapnia* sp. (26 control and 27 treatment), 8 aquifer *P. frontalis* (4 control and 4 treatment), and 8 benthic *Sweltsa* sp. (4 control and 4 treatment).

Reads were then trimmed and filtered based on quality and length using Trimmomatic (44) and FastQC v0.12.1 (45). Adapters were removed from reads, and a 4-base sliding window was used to trim read ends. The minimum allowable read length was 60 bp.

### De Novo Assembly

*De novo* transcriptome assembly was conducted on filtered reads for each sample using the Trinity assembler version 2.2.1 (46). Redundant and extremely low-expressed contigs (consensus regions from overlapping reads) were removed using the filter_fasta_by_resem_values.pl Trinity-utility. A minimum contig length of 300 nucleotides was used, and default in silico normalization was applied with a target maximum coverage of 50 reads. Parameters included a minimum k-mer coverage of 2, an expected fragment length of 500 bp, and assembly using the original Trinity algorithm (Inchworm, Chrysalis, and Butterfly). A 98% identity threshold was maintained for path merging, allowing a maximum of two mismatches and a maximum internal gap of 10 bp. In addition, supertranscripts were generated to represent alternative isoforms within each locus.

Assembly was evaluated using BUSCO v6 (47) in transcriptome mode with the phylogenetic lineage arthropoda_odb10 (1,013 orthologs), applying the default search threshold and an e-value of 1 × 10⁻³. The analysis used the complete set of assembled transcripts (*Isocapnia, or Paraperla or Sweltsa*, separately) as input and reported the standard BUSCO categories: complete orthologs (single-copy or duplicated), fragmented, and missing.

To reduce redundancy in the transcript assembly, CD-HIT-EST (48) was applied using a 95% identity threshold in global mode. Default parameters were used for sequence comparison, including a minimum alignment length of 10 nucleotides, a bandwidth of 20, and a word size of 10. Coding region prediction was performed using TransDecoder v5.5.0 (https://github.com/TransDecoder/TransDecoder) based on the dataset reduced with CD-HIT at 95% identity. The universal genetic code was used, with a minimum protein length of 100 amino acids, assuming non–strand-specific data (universal genetic code, minimum protein length = 100 aa, strand-specific = false, Pfam search = true, Single Best Only = false, 500 long ORFs used for training).

### Functional annotation

Functional annotation of the predicted proteins was performed using a combined approach. First, DIAMOND v2.1.8 (49) was run in BLASTp mode against the NCBI NR database, using the following parameters: e-value ≤ 1e-3, more-sensitive mode, maximum number of retrieved hits = 20, minimum HSP length = 33, and no initial coverage restriction (HSP-hit coverage = 0). Three alternative taxonomic filters were applied in separate runs (Arthropoda, Hexapoda, Insecta) to prioritize biologically relevant alignments. BLAST results were processed using Blast2GO version 5.2.5 (50), mapping hits to Gene Ontology (GO) terms. The annotation rule was then applied to assign GO terms to each protein. Annotation was further complemented with eggNOG-mapper v2.1.0 (51), using the eggNOG 5.0.2 database, to enrich the assignment of orthologous functions, COG categories, and KEGG metabolic pathways. Finally, Blast2GO and eggNOG annotations were integrated, removing redundant terms per sequence.

Abundance quantification was performed using the Transcript-level Quantification tool based on the RSEM package (52). This method assigns multi-mapped reads to genes and isoforms using an expectation–maximization approach, providing robust estimates at both the transcript and gene levels. The paired-end libraries were processed under a non–strand-specific scheme, with estimation of the read start position distribution (Estimate RSPD = true) and without the addition of artificial poly(A) tails.

### Differential expression analysis

Differential expression analysis was conducted using the edgeR package (version 3.10; 53) in R (version 3.3; 54). Before analysis, an automatic low-expression filtering step based on CPM (counts per million) was applied, retaining only genes with sufficient counts in a minimum number of samples. Data were normalized using the TMM (Trimmed Mean of M-values) method, which corrects for differences in sequencing depth and library composition. Differentially expressed genes were identified based on a false discovery rate (FDR) < 0.01 and an absolute log₂ fold change > 1. Pairwise differential gene expression (DEG) analysis was performed between control and hypoxia-treated samples using the glm function in R. The Benjamini-Hochberg false discovery rate of 1% was applied using the p.adjust function. Statistical inference was performed using a Generalized Linear Model (GLM) with a robust Likelihood Ratio Test (LRT), thereby improving stability in the face of atypical dispersion estimates.

Given the temporal nature of the experimental design (0, 10, and 60 min) for *Isocapnia* sp., differential expression analysis was performed using the maSigPro package (55) in R, which is designed to detect dynamic changes in time-series experiments. Briefly, the raw counts were first normalized using the TMM method in edgeR. Then, a quadratic polynomial regression approach (degree = 2) was applied. This polynomial degree was selected to capture nonlinear trends and to enable the detection of complex transient responses. Global significance was evaluated using the *p.vector* function with a Benjamini–Hochberg correction (FDR < 0.05). To refine the selection of significant genes, stepwise regression was applied using the *two.ways.forward* method in R. This configuration is robust for multifactorial designs with batch correction, allowing specific identification of genes whose significant changes are driven by the Time–Treatment interaction, while excluding those responding only to technical effects (56). A strict goodness-of-fit filter was applied, retaining only genes with R² > 0.6, ensuring that the model explains at least 60% of the observed variance. Finally, the resulting DEGs were grouped into nine clusters using hierarchical clustering in R to facilitate interpretation of co-regulated expression patterns. These clusters represent groups of genes sharing similar expression trajectories under treatment relative to control conditions, indicating coordinated regulatory responses to environmental stress.

### Enrichment and metabolic routes

To assign biological meaning to the identified temporal expression patterns, an Over-Representation Analysis (ORA) was performed independently for each of the nine clusters generated by maSigPro for *Isocapnia* sp using the clusterProfiler package in R (57). The objective was to identify which biological processes, molecular functions, or cellular components are significantly enriched within each group of co-regulated genes.

Functional enrichment of differentially expressed transcripts was evaluated using one-tailed Fisher’s exact test, applied independently to the sets of up-regulated and down-regulated genes (*P*. *frontalis* and *Sweltsa* sp.) or genes within a cluster (*Isocapnia* sp.). The three main categories of GO were included: Biological Process (BP), Molecular Function (MF), and Cellular Component (CC). The test was applied with multiple-testing correction using the Benjamini–Hochberg method (FDR), with a significance threshold of adjusted p < 0.05, in Blast2GO. Results were visualized using bubble plots that integrated statistical significance (color) and gene count (bubble size), facilitating rapid identification of key biological functions activated or suppressed by hypoxia.

Gene Set Enrichment Analysis (GSEA) was applied to identify coordinated expression patterns in predefined gene sets based on GO terms using Blast2GO. Unlike approaches focused on individual genes, GSEA assesses whether a set of related genes exhibits significant enrichment at the top or bottom of the list of differentially expressed genes (DEGs). For this analysis, genes were ranked using a value calculated as:

Rank = sign(logFC) × −log10(P-value)

The analysis was performed with 1,000 permutations, using a weighted enrichment statistic, and included the three main GO categories (Biological Process, Molecular Function, Cellular Component). Gene set size limits were set between 15 and 500, and an FDR < 0.25 was applied, following the standard criterion recommended for GSEA.

Finally, pathway enrichment analysis was conducted using the KEGG database via Blast2GO, integrating information from GO functional annotations, enzyme codes (EC), and annotated sequences. KEGG provides manually curated maps of molecular interactions, metabolic pathways, and cellular processes, organized into orthologous groups (KO). All analyses were performed with 1,000 permutations, a weighted enrichment statistic, gene set size limits of 15 (minimum) and 500 (maximum), and an FDR threshold of ≤0.25. Additionally, a complementary enrichment analysis was performed using Fisher’s exact test, with multiple comparisons corrected using the Benjamini–Hochberg method (FDR < 0.05). All results were summarized in Supplementary Table 1.

## Data, Materials and software availability

Data for this study are available at the Sequence Read Archive (SRA) database under the Bioproject accession number (xxx).

## Acknowledgments

We thank the Dalimata family for allowing us to conduct this research on their land. We thank Julia Cotter and Haley Dole for assistance with sample collection and laboratory experiments. This work was funded by a National Science Foundation Dimensions of Biodiversity Grant #DOB-1639014. The computational analysis was supported under the National Laboratory for High Performance Computing (NLHPC) research project category by the Universidad de Chile. This research was supported by Fondecyt regular grant #1240712 from the Agencia Nacional de Investigación y Desarrollo (ANID) to MG.

## Author contribution

M.G. conceived the analysis, wrote the code, analyzed data, and drafted the manuscript. M.V. analyzed Isocapnia data. E.W. laboratory experiments, writing, editing. B.H. funding, project conceptualization, editing, and data cleaning. G. L. funding, project conceptualization, editing. J. S. funding, project conceptualization, editing. R. M. project conceptualization, laboratory experiments, writing, and editing.

## Supplementary files

**Supplementary Table 1.** Summary of read/transcript reduction and dataset usage across analysis.

| Step | *Isocapnia*<br>sp. | *Paraperla*<br>*frontalis* | *Sweltsa* sp. |
| --- | --- | --- | --- |
| Raw reads<br>(initial) | ~1 billion<br>reads total | ~200 million<br>reads total | ~300 million<br>reads total |
| Quality filtering /<br>trimming | 255,157,564<br>reads | 132,364,034<br>reads | 216,958,202<br>reads |
| Initial<br>transcriptome<br>(Trinity assembly) | 262,946<br>transcripts | 126,900<br>transcripts | 193,260<br>transcripts |
| Post-clustering<br>(CD-HIT, 95%) | Reduced to<br>192,788<br>transcripts | Reduced to<br>100,762<br>transcripts | Reduced to<br>125,325<br>transcripts |
| BUSCO | 97.43% | 97.63% | 98.52% |
| Clustering | 192,788<br>clusters | 100,762<br>clusters | 125,325<br>clusters |
| Coding sequence<br>prediction<br>(TransDecoder) | 48,583 ORFs | 35,491 ORFs | 37,406 ORFs |
| Functional<br>annotation | 48,582<br>proteins | 35,491<br>proteins | 37,406<br>proteins |
| Differential<br>expression (total<br>genes) | 6,004 | 1,762 | 1,298 |

**Supplementary Table 2.** Functional annotations of selected genes among cluster and their isoforms obtained in *Isocapnia* sp. GO, gene ontology.

| GO Term | GO Name | Functional Annotation | P-value | Isoforms |
| --- | --- | --- | --- | --- |
| GO:0006119 | oxidative phosphorylation | Energy production (mitochondrial respiration) | ~0.0028–0.0032 | 2–6 |
| GO:0000096 | sulfur relay system | Sulfur metabolism / cofactor synthesis | ~0.003 | 2–6 |
| GO:0006772 | thiamine metabolism | Cofactor metabolism | ~0.002–0.008 | 10–25 |
| GO:0006163 | purine metabolism | Nucleotide metabolism | ~0.002–0.008 | 10–25 |
| GO:0042254 | ribosome biogenesis | Protein synthesis machinery | ~0.004–0.008 | 10–25 |
| GO:0032956 | regulation of actin cytoskeleton | Cell structure and movement | ~0.003–0.005 | 5–7 |
| GO:0006909 | phagocytosis | Immune / cellular uptake | ~0.003–0.005 | 5–7 |
| GO:0006526 | arginine and proline metabolism | Amino acid metabolism | ~0.005–0.008 | 5–8 |
| GO:0006558 | phenylalanine metabolism | Amino acid metabolism | ~0.005–0.008 | 5–8 |
| GO:0000165 | MAPK signaling pathway | Stress and signaling pathways | ~0.005–0.008 | 5–8 |
| GO:0034976 | protein processing in ER | Protein folding / stress response | ~0.002–0.007 | 4–7 |
| GO:0007179 | TGF-beta signaling pathway | Cell signaling / regulation | ~0.002–0.007 | 4–7 |
| GO:0009396 | folate biosynthesis | Cofactor metabolism | ~0.002–0.007 | 4–7 |
| GO:0006749 | glutathione metabolism | Oxidative stress response | ~0.001–0.005 | 4–16 |
| GO:0005840 | ribosome | Protein synthesis | ~0.001–0.005 | 4–16 |
| GO:0019439 | dioxin degradation | Detoxification pathways | ~0.001–0.005 | 4–16 |
| GO:0031103 | axon regeneration | Neural repair / signaling | ~0.0014 | 3 |
| GO:0007214 | GABAergic synapse | Neural signaling | ~0.002–0.006 | 2–5 |
| GO:0019752 | butanoate metabolism | Energy metabolism | ~0.002–0.006 | 2–5 |
| GO:0005777 | peroxisome | Oxidative metabolism / detox | ~0.002–0.006 | 2–5 |
| GO:0016050 | primary bile acid biosynthesis | Lipid metabolism | ~0.002–0.006 | 2–5 |
| GO:0006098 | pentose<br>phosphate<br>pathway | Redox / NADPH<br>production | ~0.002–<br>0.006 | 5–12 |
| GO:0007186 | neuroactive<br>ligand-receptor<br>interaction | Cell<br>communication | ~0.002–<br>0.006 | 5–12 |
| GO:0006941 | vascular smooth<br>muscle<br>contraction | Muscle /<br>structural<br>function | ~0.002–<br>0.006 | 5–12 |

**Supplementary Table 3.** Functional annotations of selected genes and their isoforms obtained in *P. frontalis* GO, gene ontology.

| GO Term | GO Name | Functional Annotation | p-value | Isoforms |
| --- | --- | --- | --- | --- |
| GO:0005576 | extracellular region | Cellular Component | 4.69E-50 | 136 |
| GO:0110165 | cellular anatomical structure | Cellular Component | 1.00E-35 | 691 |
| GO:0032501 | multicellular organismal process | Biological Process | 8.52E-30 | 328 |
| GO:0071944 | cell periphery | Cellular Component | 1.54E-29 | 188 |
| GO:0016020 | membrane | Cellular Component | 5.45E-29 | 315 |
| GO:0005615 | extracellular space | Cellular Component | 1.46E-27 | 72 |
| GO:0009987 | cellular process | Biological Process | 4.91E-27 | 643 |
| GO:0050896 | response to stimulus | Biological Process | 1.31E-24 | 294 |
| GO:0009605 | response to external stimulus | Biological Process | 1.37E-24 | 143 |
| GO:0048856 | anatomical structure development | Biological Process | 2.18E-22 | 275 |
| GO:0007275 | multicellular<br>organism<br>development | Biological<br>Process | 2.17E-22 | 250 |
| GO:0005886 | plasma<br>membrane | Cellular<br>Component | 6.94E-22 | 151 |
| GO:0003824 | catalytic activity | Molecular<br>Function | 6.49E-22 | 357 |
| GO:0032502 | developmental<br>process | Biological<br>Process | 7.53E-21 | 280 |
| GO:0042221 | response to<br>chemical | Biological<br>Process | 4.13E-17 | 144 |
| GO:0005488 | binding | Molecular<br>Function | 2.28E-16 | 429 |
| GO:0043167 | ion binding | Molecular<br>Function | 2.92E-16 | 218 |
| GO:0009653 | anatomical<br>structure<br>morphogenesis | Biological<br>Process | 3.92E-16 | 178 |
| GO:0048731 | system<br>development | Biological<br>Process | 4.79E-16 | 193 |
| GO:0008037 | cell recognition | Biological<br>Process | 6.04E-15 | 42 |

**Supplementary Table 4.**
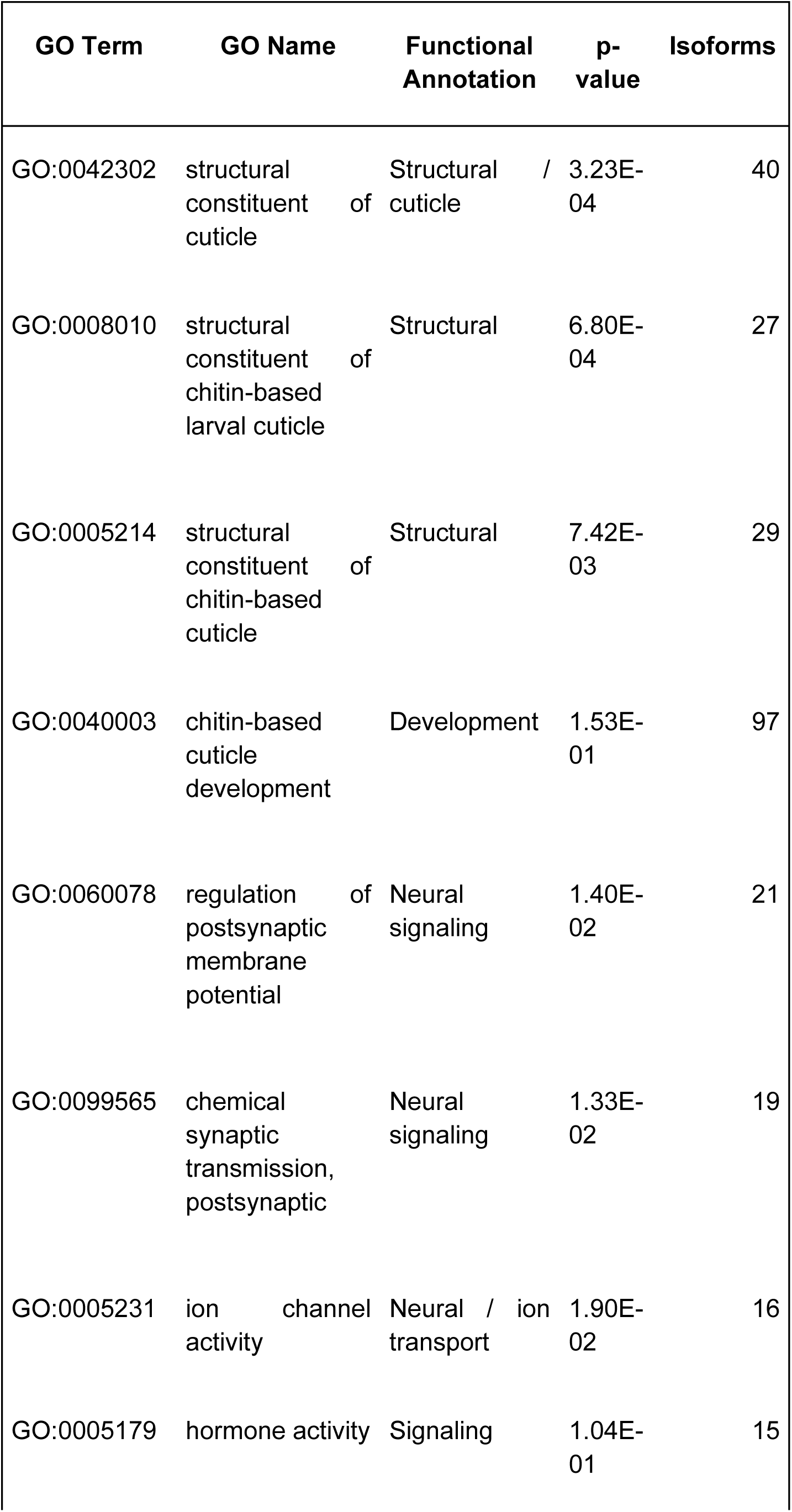
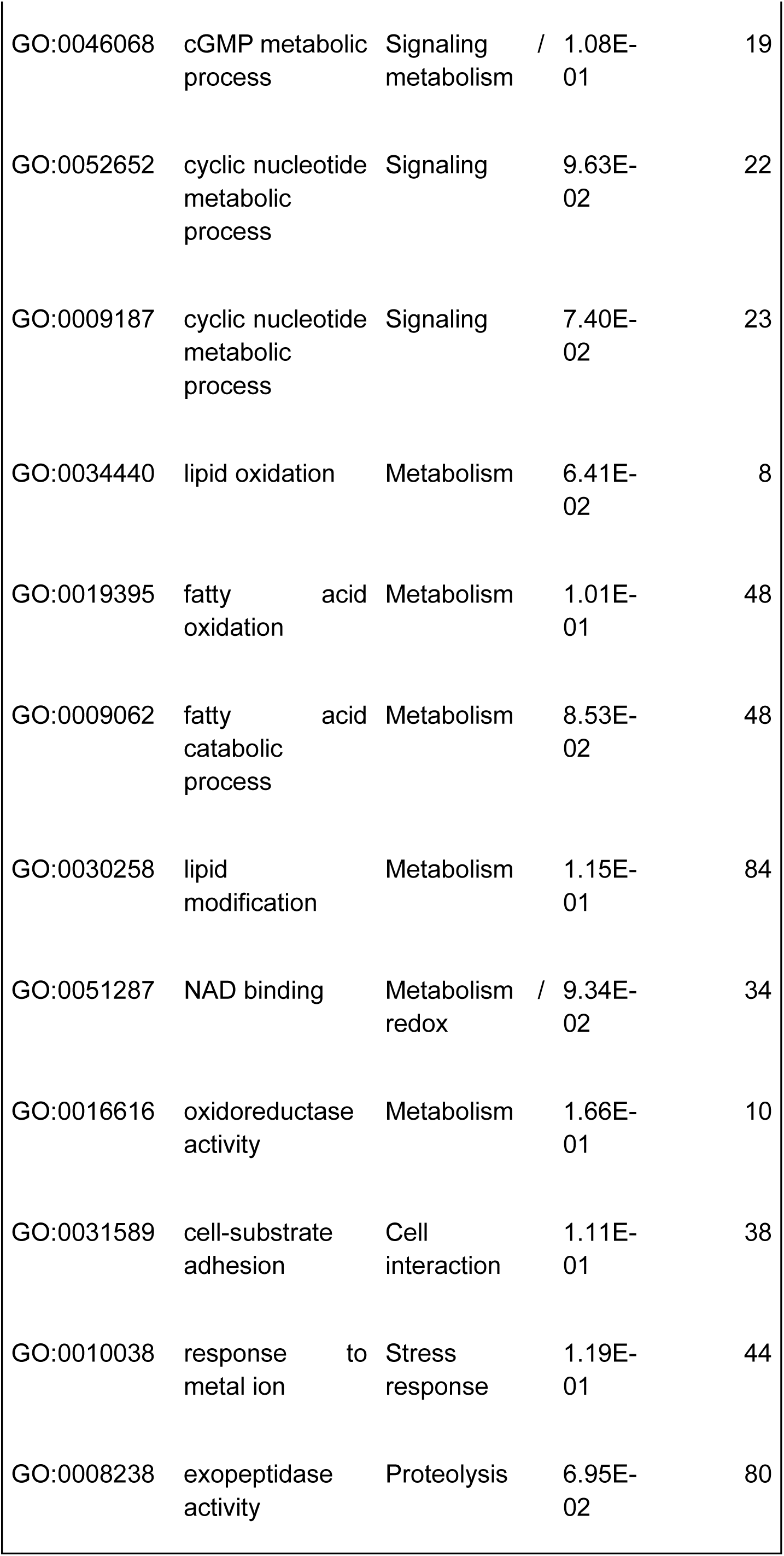
Functional annotations of selected genes and their isoforms obtained in *Sweltsa* sp. GO, gene ontology.

**Supplementary Figure 1.**
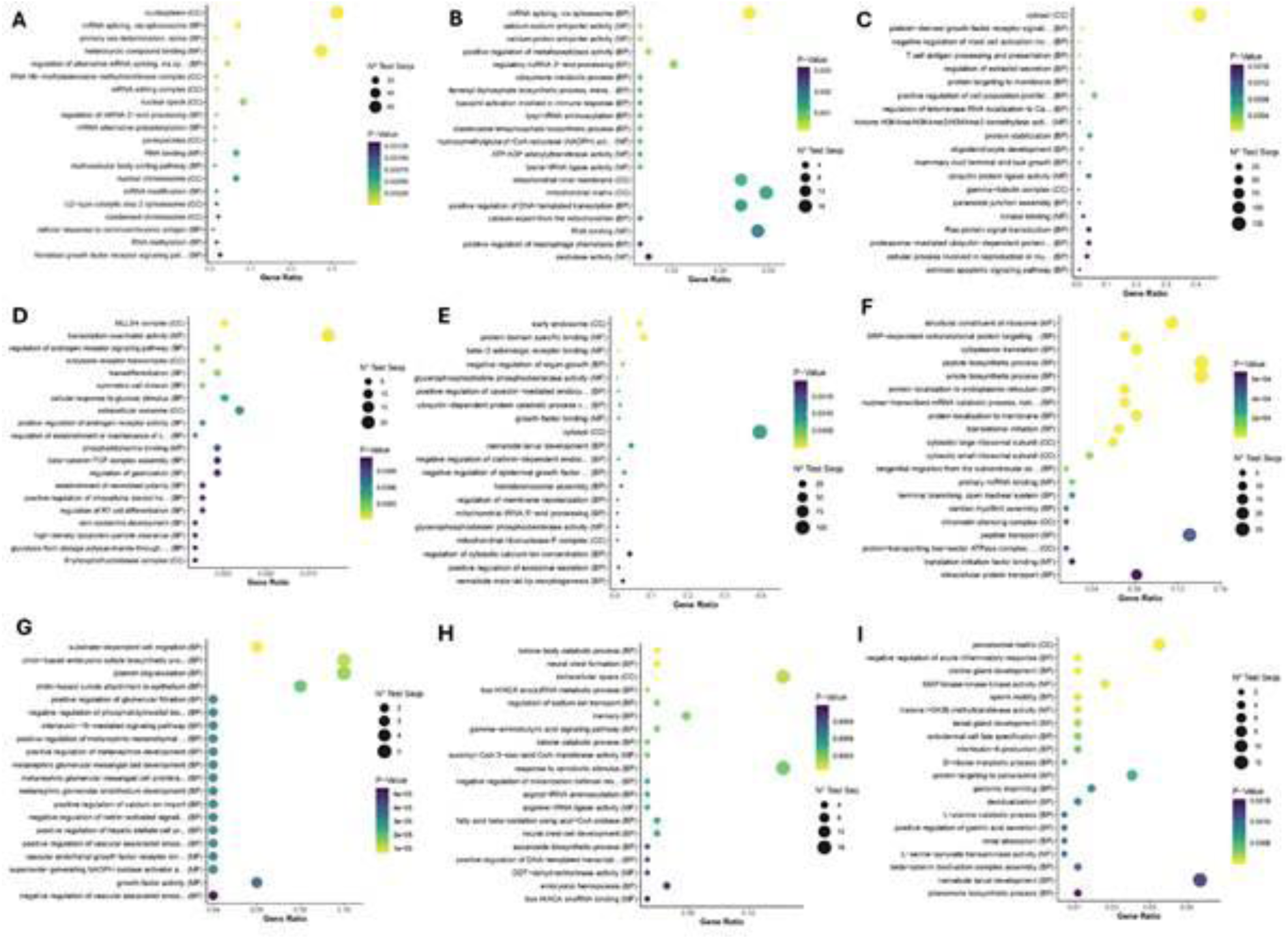
Gene ontology enrichment across experimental conditions. (A–I) Enriched Gene Ontology (GO) categories from differentially expressed genes across 9 clusters for *Isocapnia* sp. Rows correspond to conditions and columns to functional classifications. The x-axis shows gene ratio, point size indicates gene count, and color represents adjusted *P* values. Only top 20 are shown.

**Supplementary Figure 2.**
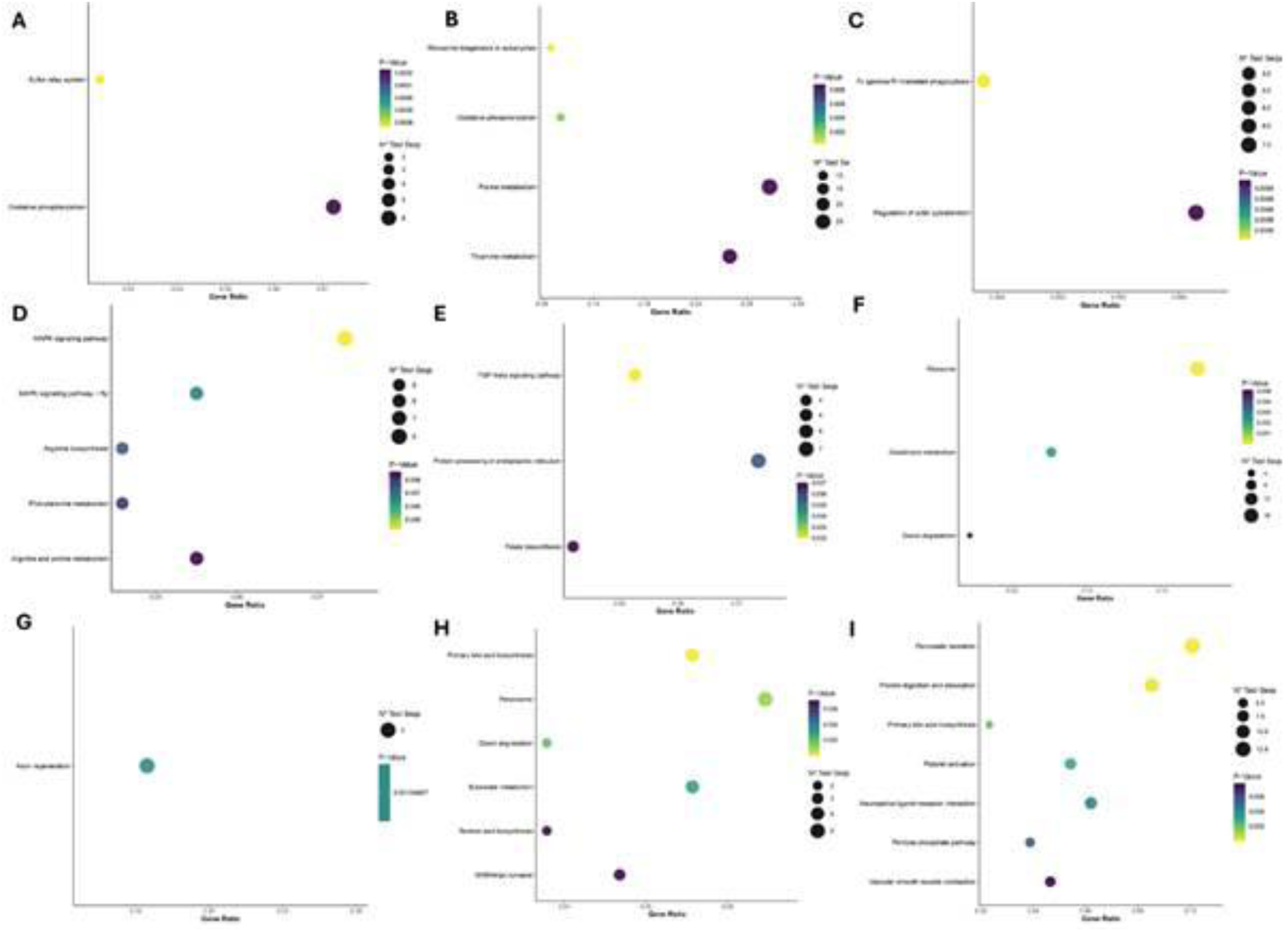
Kyoto Encyclopedia of Genes and Genome (KEGG) pathway enrichment across experimental conditions for *Isocapnia* sp. (A–I) Enriched KEGG pathways identified from differentially expressed genes across 9 clusters. Rows correspond to conditions and columns to distinct comparisons or gene sets. The x-axis shows gene ratio, point size indicates the number of genes per pathway, and color represents adjusted *P* values. Only top 20 are shown.

**Supplementary Figure 3.**
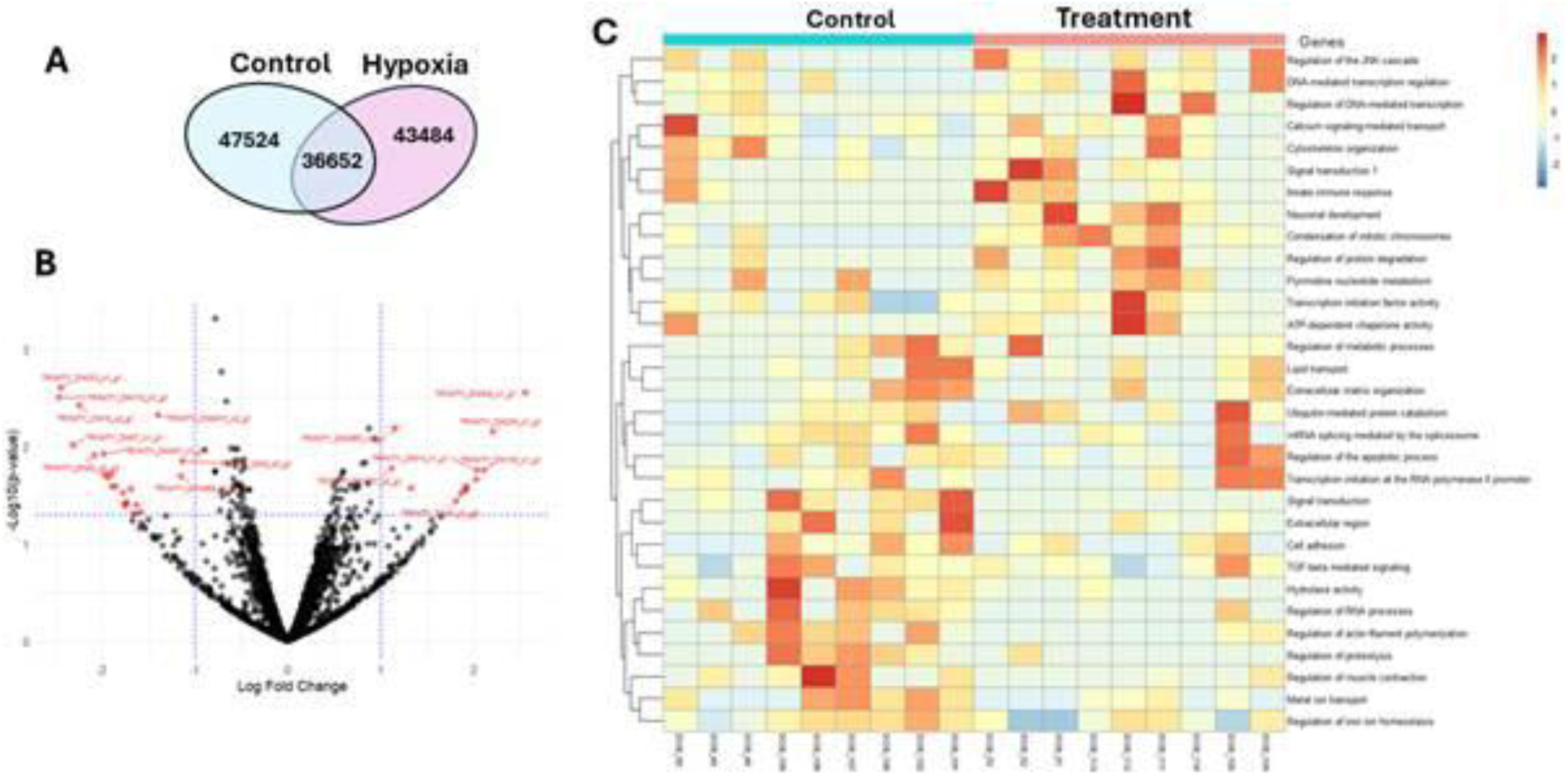
Differential expression analysis for *Isococania* sp. (A) Venn diagram of the total number of orthologous genes detected in control and hypoxia treatments. (B) Volcano plot showing differential expression results, with significantly regulated genes (red) at FDR < 0.05; grey points indicate non-significant genes. (C) Heatmap of 32 differentially expressed genes annotated by function between control and hypoxia treatments. Rows represent genes, grouped and labeled according to their functional annotation, and columns represent individual samples ordered by treatment. Gene expression values are scaled by row, with warmer colors indicating higher relative expression and cooler colors indicating lower expression. Hierarchical clustering of samples shows separation between control and hypoxia groups, while clustering of genes reveals functionally related sets with similar expression patterns across treatments.

**Supplementary Figure 4.**
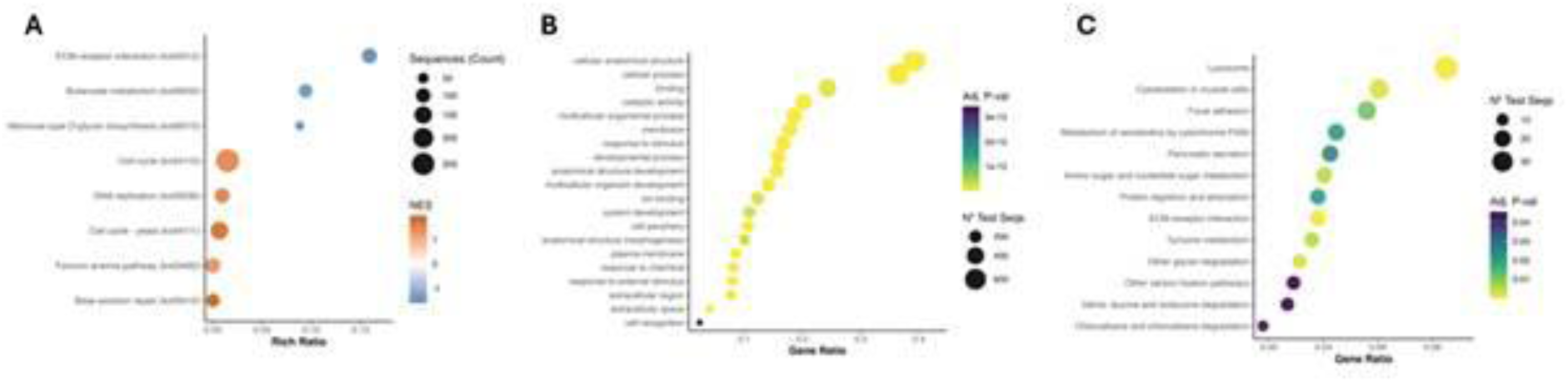
Functional enrichment of up- and downregulated genes for *Paraperla frontalis*. (A) Top 20 enriched KEGG pathways from differentially expressed genes. (B) Top 20 Gene Ontology (GO) terms enriched among upregulated genes based on Fisher’s exact test. (C) Top 20 Gene Ontology (GO) terms enriched among downregulated genes based on Fisher’s exact test. The x-axis shows gene ratio, point size indicates gene count, and color represents adjusted *P* values.

**Supplementary Figure 5.**
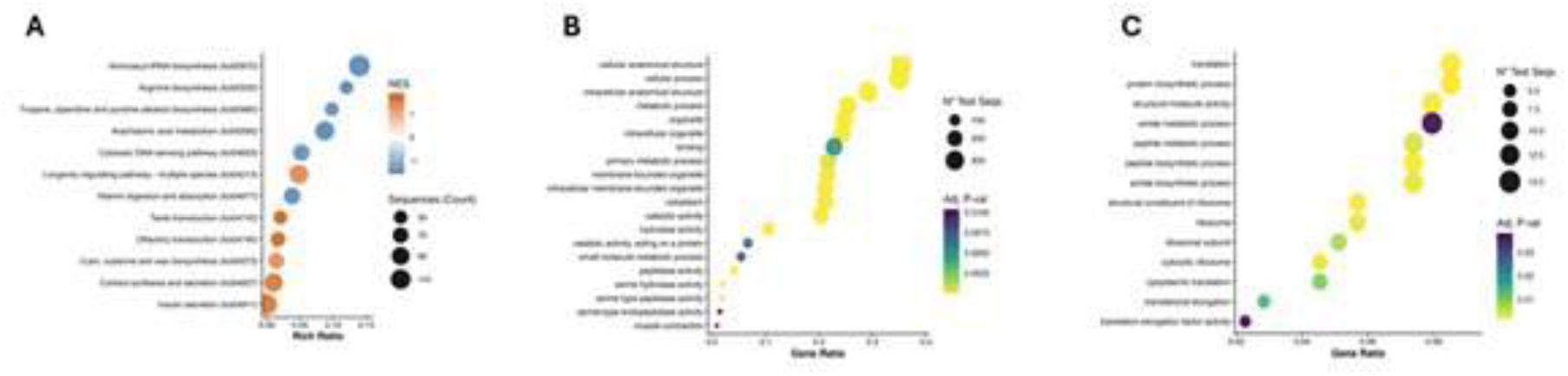
Functional enrichment of up- and downregulated genes for *Sweltsa* sp. (A) Top 20 enriched KEGG pathways from differentially expressed genes. (B) Top 20 Gene Ontology (GO) terms enriched among upregulated genes based on Fisher’s exact test. (C) Top 20 Gene Ontology (GO) terms enriched among downregulated genes based on Fisher’s exact test. The x-axis shows gene ratio, point size indicates gene count, and color represents adjusted *P* values.

**Supplementary Figure 6.**
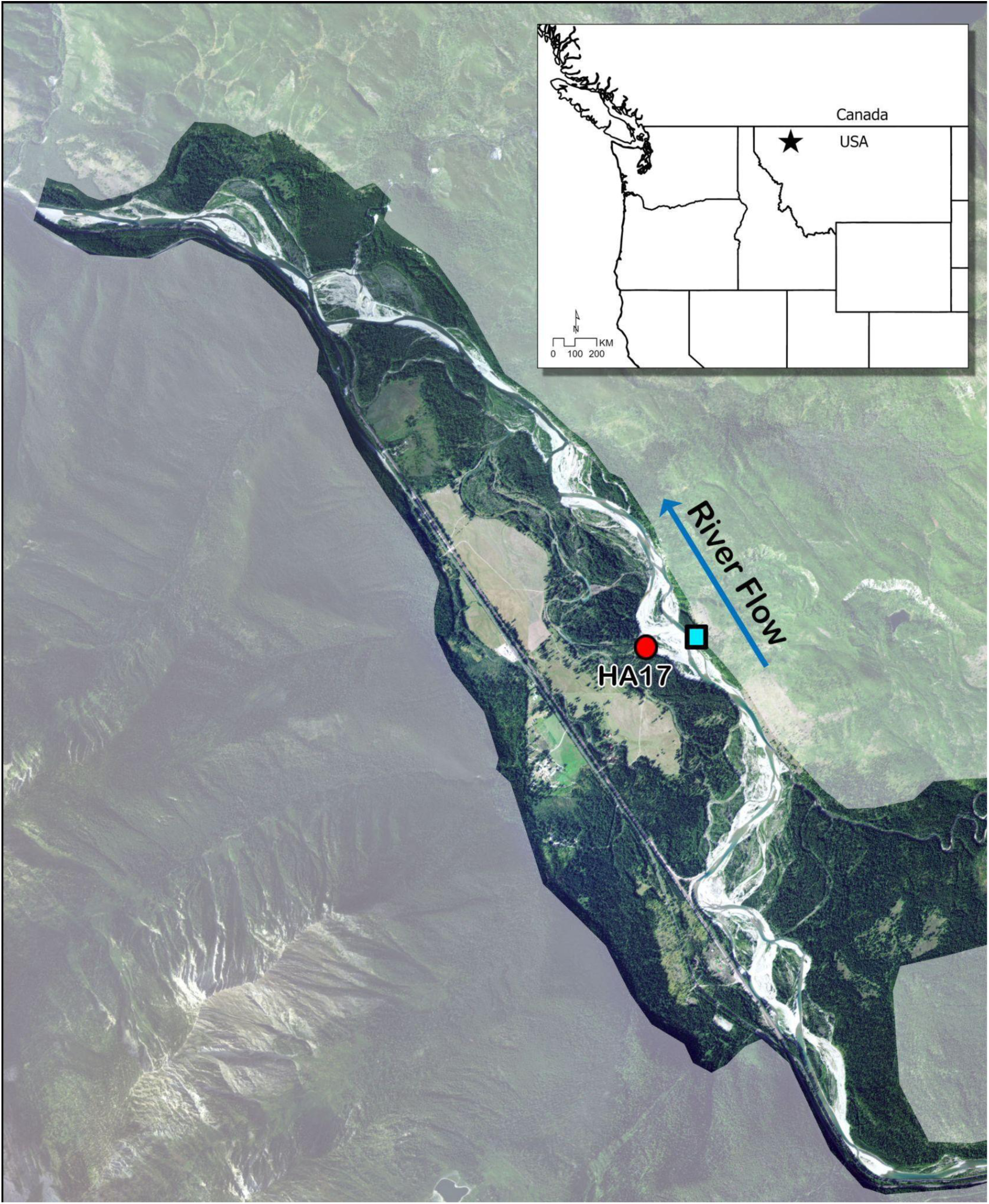
Stonefly nymphs were collected from aquifer (red circle) and benthic habitats (blue square) the Nyack floodplain in NW Montana.

